# Differentiating the mechanism of antibacterial activities of nano and ionic copper by using *Escherichia coli* as a model microorganism

**DOI:** 10.1101/2024.10.05.616784

**Authors:** Gregor P. Jose, Subhankar Santra, Saurav Kumar Saha, Swadhin K Mandal, Tapas K. Sengupta

## Abstract

In this study, the effect of polymer stabilized copper nanoparticles and ionic copper on the growth, nucleic acid pool, reactive oxygen species generation, cell surface lipopolysaccharide, outer membrane protein profile and cell surface morphology of *Escherichia coli* were investigated. Copper nanoparticles exhibited a superior bactericidal activity associated with increased nucleic acid degradation, reactive oxygen species generation and change in the outer membrane protein profile compared to ionic copper in a concentration dependent manner. Although, there was no change in the outer membrane lipopolysaccharide profile, inductively coupled plasma mass spectrometry analysis of nano- and ionic copper treated *Escherichia coli* cells revealed that more amounts of copper nanoparticles were transported inside the cells compared to the ionic counterpart up to 500 μM concentrations. Interestingly, copper nanoparticles at 1000 μM concentration could induce membrane pit formation whereas ionic copper failed to exhibit such property under the same experimental conditions. Based on these observations it can be concluded that both nano- and ionic copper exert their antibacterial action through the generation of reactive oxygen species, degradation of cellular nucleic acids and alteration of membrane protein profile, but with a significant difference in the effective concentration range due to the differential cellular transport.

## 1. Introduction

Copper is an essential micronutrient for the bacteria. Its major function is to act as a cofactor for different metalloenzymes including copper nitrite reductase, nitric oxide reductase, nitrous oxide reductase involved in the denitrification process, bacterial cytochrome C oxidase associated with the generation of the electrochemical gradient for the ATP production, tyrosinase enzyme involved in the melanin production etc (Fairhead and Thöny-Meyer 2012; Kaila et al. 2009; Tavares et al. 2006; Tsukihara et al. 1995). But free ionic copper, due to its high reactivity can generate highly reactive derivatives like hydroxyl radical, superoxide anions, and hydrogen peroxide mediated by the ‘Fenton reaction’(Cross et al. 2003; Fenton 1894; Sutton and Winterbourn 1989). This oxidative stress poses a great threat to the survival of bacteria. Ancient civilizations recognized this remarkable potential of copper and they utilized copper salts for various medicinal purposes. The first documented evidence of use of copper salts was found in two Egyptian medicinal papyri, the Ebers papyrus and Edwin Smith’s surgical papyrus. They were regarded as the oldest surviving medical manuscripts in the world (Hostynek and Maibach 2006). The Ebers papyrus lists different forms of copper including copper rust, copper vitriol, copper verdigris etc, under the mineral remedies section (Bryan and Joachim 1930). Edwin Smith’s surgical papyrus describes the use of copper for treating infected chest wounds (Grass et al. 2011). In 400 BC, Hippocrates described the detailed medicinal use of copper to treat leg ulcers (O’Gorman and Humphreys 2012). In India, Charaka Samhita (300-500 AD) details the use of Tamra (copper) formulations for treating tuberculosis (*ksaya*), obesity (*sthaulya*), leprosy (*kusta*), respiratory diseases (*kasa*) etc (Galib et al. 2011). Most remarkably, during the 19th-century cholera epidemic in Paris, copper workers were found immune to the disease (Grass et al. 2011) suggesting the role of copper in resisting the disease.

Although copper had a glorious past, the development of antibiotics in the 19^th^ century diminished the use of metallic copper for clinical applications. The development of sulphonamides, penicillin, and aminoglycosides has completely revolutionized the treatment regime and saved millions of patients from death (Powers 2004). During the last few decades, the rise of many antibiotic-resistant strains of bacteria posed a great challenge to both clinicians and researchers (Andersson and Hughes 2010; Neu 1992). Nosocomial infections or Health Care Associated Infections (HCAI) caused by methicillin-resistant *Staphylococcus aureus* (MRSA) and vancomycin-resistant enterococci (VRE) showed a rapid increase in the latter half of the 20^th^ century, mainly in the developing countries (Hsueh et al. 2005; Okeke et al. 2005). This situation rekindled the interest in the use of metallic copper surfaces or alloys to combat nosocomial infections. Recent studies showed a very promising action of these metallic copper surfaces against common hospital born infectious agents like MRSA, VRE, *Clostridum difficile* etc (Michels et al. 2009; Schmidt et al. 2012; Warnes and Keevil 2011; Wheeldon et al. 2008, Aillón-García et al. 2023). This bactericidal action of metallic copper is known as “contact killing”(Grass et al. 2011). The major participants in the “contact killing” process are the copper ions released from the interacting zone between bacteria and copper surface. This mechanism is always under threat by the evolution of ionic copper-resistant bacteria which can form biofilm on copper surfaces (Kielemoes and Verstraete 2001).

To prepare for this alarming possibility, we need alternate forms of copper with superior bactericidal activities and the copper nanoparticles (CuNPs) fill this gap perfectly. The change in its size from a bulk level to the nanoscale level has imparted different unique optical and catalytic properties to the CuNPs like distinctive distribution, large surface to volume ratio resulting in enhancement of its application (Kelly et al. 2002; Narayanan and El-Sayed 2005, Xu et al. 2022). The incorporation of CuNPs into materials like soda lime glass, sepiolite, chitosan and polyelectrolyte has generated materials with potent bactericidal properties (Esteban-Cubillo et al. 2006; Esteban-Tejeda et al. 2009; Mallick et al. 2012, Jessop et al. 2021). Antibacterial cotton, containing CuNPs were found to be effective against nosocomial infections caused by *Acinetobacter baummanii* (Cady et al. 2011). There are a number of studies present in the literature that showed CuNPs induced antibacterial as well as antiviral action (Ramos-Zuniga et al. 2023, Ahmadi et al. 2023). Yoon *et al* compared the antibacterial action of both silver and CuNPs against *Escherichia coli* and *Bacillus subtilis*. They found that CuNPs exhibited a superior bactericidal potential than the silver nanoparticles (Yoon et al. 2007). Ruparelia *et al* studied the antibacterial activity of CuNPs against *Escherichia coli, Bacillus subtilis*, and *Staphylococcus aureus* cells, and speculated that the antibacterial activity of CuNPs was mediated by the release of ionic copper inside the cells, leading to the disruption of biochemical processes and distortion of the DNA helical structure (Ruparelia et al. 2008). Pramanik *et al* also reported the antibacterial mechanism of copper iodide (CuI) nanoparticles against *Escherichia coli* and *Bacillus subtilis* cells (Pramanik et al. 2012). Recent studies showed that CuO nanoparticles could exhibit significant activity against the biofilm formation of MRSA and some Gram-negative bacteria (Agarwala et al. 2014, Bai et al. 2022). Qamar et al. recently showed the potential of copper nanoparticles against various multi-drug resistant bacterial strains (Qamar et al, 2020). CuO nanoparticles also showed a marked reduction in the human oral biofilm load by inhibiting the growth of oral bacteria (Tabrez Khan et al. 2013). The most probable mechanism behind the CuNPs toxicity is by means of “Trojan Horse” mode, which allows metal nanoparticles to cross through biological membranes and transport metal cargo in to the cells (Studer et al. 2010). However, there are no comprehensive studies available in the literature addressing the difference in the mechanism of antibacterial action of nano and ionic copper.

Previous studies from our laboratory revealed the difference in the abilities of CuNPs and ionic copper to degrade DNA *in vitro*. It was observed that CuNPs can degrade isolated DNA molecules *in vitro* in a dose dependent manner mediated through the generation of singlet oxygen. On the other hand, ionic copper did not show any such *in vitro* DNA degrading activity(Jose et al. 2011). In the present study we have compared the mechanism of the antibacterial effect of CuNPs and copper sulfate (Cu^2+^) on *Escherichia coli*. The results point towards a better bactericidal activity shown by CuNPs over their ionic counterparts due to their nano size and better transport into the cells.

## 2. Materials and Methods

### 2.1 Preparation and characterization of copper nanoparticles

#### 2.1.1 Synthesis of copper nanoparticles

CuSO_4_.5H_2_O (0.0624 g, 0.25 mmol) and poly (4-styrene-sulfonic acid-*co*-maleic acid) sodium salt (average M_w_ = 20,000; 60 mg, 3 μmol) were taken in a 50 mL Schlenk flask followed by the addition of 10 mL deionized water and stirred at room temperature (25 °C). In a 25 mL Schlenk tube, 19 mg of NaBH_4_ (0.5 mmol) was taken followed by the addition of 5 mL of deionized water. Then both the solutions were degassed under a high vacuum followed by filling up with N_2_ using the standard Schlenk line technique (Shriver and Drezdzon 1982). This process was repeated for 5 to 6 times to ensure complete removal of dissolved oxygen from the solutions. Then 0.5 mL of NaBH_4_ solution was added dropwise to the CuSO_4_-polymer mixture solution at room temperature under vigorous stirring. After complete addition, stirring was stopped immediately and the color of the mixture-solution was changed from blue to green and ultimately dark brown indicating the formation of CuNPs. This nanoparticle suspension was filtered using 0.22-micron filter to ensure sterility.

#### 2.1.2 Transmission Electron Microscopic (TEM) characterization

CuNP solution was characterized by placing nanoparticle suspension onto carbon coated copper grid and allowed to dry overnight. On the next day, these grids were analyzed by TEM (JEOL, JEM 2010) with an accelerating voltage of 200 kV.

### 2.2 Comparison of antibacterial action of CuNPs and CuSO_4_

*Escherichia coli* (ATCC 25922) cells (10^8^ CFU/mL) in nutrient broth were treated with various concentrations of CuNPs and CuSO_4_ (250, 500 and 1000 μM) solutions for 4 hours at 37 °C with continuous shaking at 150 rpm. A set of tubes without any added chemicals and another set with a mixture of sodium borohydride and polymer were taken as controls. After the treatment, the bacterial cultures were serially diluted (up to 10^6^ folds) using nutrient media. 100 µL of this diluted culture for each diluted sample was spread on to nutrient agar plates and incubated at 37°C overnight. On the next day, colony forming units (CFU) were enumerated. Each data point corresponds to an average of 4 independent experiments.

### 2.3 Isolation of nucleic acids from *Escherichia coli* after treatment with CuNPs and CuSO_4_

*Escherichia coli* cells (10^8^ CFU/mL) in nutrient broth were treated with 500, 750, 1000, and 1500 µM of CuNPs and CuSO_4_ separately for 4 hours. After the treatment, cells were washed with phosphate buffered saline (PBS) and re-suspended in 100 µL of distilled water. 200 µL of saturated phenol-chloroform mixture was added in to it and vortexed for 1 minute at room temperature. This was followed by centrifugation at 13200 rpm for 10 minutes at 4°C. 10 µL of the aqueous phase was separated in 1% agarose gel electrophoresis and nucleic acids were visualized in gel by staining with Ethidium bromide.

### 2.4 Reactive Oxygen Species (ROS) detection in *Escherichia coli*

The ability of CuNPs and CuSO_4_ to induce the production of reactive oxygen species (ROS) in bacterial cells was analyzed by using 2’,7’-dichlorodihydrofluorescein diacetate (H_2_DCFDA, Invitrogen) as per the manufacturer’s instruction. The non-fluorescent H_2_DCFDA is converted to the highly fluorescent 2’,7’-dichlorofluorescein (DCF) on interacting with ROS generated inside the bacterial cells and serves as an indicator for oxidative stress. In this experiment, *Escherichia coli* cells (10^8^ CFU/ mL) were treated with 1000 µM CuNPs and CuSO_4_. *Escherichia coli* cells without any treatment served as the negative control and cells with 200 µM H_2_O_2_ as the positive control. After the treatment, the cells were washed with 1 mL of saline and re-suspended in 1 mL of saline with 20 µM H_2_DCFDA followed by incubation in the dark for 1 hour. The samples were then washed with 1 mL saline and re-suspended in 100 µL of saline. 10 µL of the re-suspended sample was loaded onto glass slides and visualized using an epifluorescence microscope (Olympus-IX 81 equipped with Hamamatsu camera) with a blue excitation filter (460-495 nm) and emission filter (510-550 nm) (Gunawan et al. 2011). Image analysis was carried out using “Image Pro 6.3 software”.

### 2.5 Scanning electron microscopy (SEM)

For Scanning electron microscopy analysis, *Escherichia coli* cells were treated with 1000 µM CuNPs and CuSO_4_ for 4 hours. After the treatment, the cells were fixed with 2.5% glutaraldehyde in PBS overnight at room temperature. The cells were then washed twice with PBS and stained with 0.1% Osmium tetroxide solution for 30 minutes. The samples were washed thrice in PBS. Dehydration of the samples was carried out using graded ethanol (30, 50, 70, 90,100 % ethanol). 5 µL of the dehydrated sample was deposited onto to silicon wafer and dried overnight under a vacuum in a desiccator. SEM images were acquired by field emission scanning electron microscopy (FE-SEM, Zeiss Supra 55VP, Carl Zeiss AG, Germany) (Cho et al. 2005)

### 2.6 Surface area measurement

Surface area of the bacteria was measured from the SEM images using ImageJ software. Each data point corresponds to an average of 10 independent measurements.

### 2.7 SDS-PAGE analysis of cell surface lipopolysaccharide (LPS) profile

The LPS profile of *Escherichia coli* cells untreated and treated with either CuNPs or CuSO4 was analyzed by following the method described by Hitchcock *et al* (Hitchcock and Brown 1983). For that, *Escherichia coli* cells were treated with various concentrations of CuNPs and CuSO_4_ (250, 500, 1000 μM) solutions for 4 hours at 37°C with continuous shaking at 150 rpm. A set of tubes without any added chemicals and another set with a mixture of sodium borohydride and polymer were taken as controls. After the treatment, cells were harvested by spinning at 3341 g (6000 rpm) at 4°C for 10 minutes. The pellet was suspended in 100 μL gel lysis buffer (containing 100 mM Tris-cl pH 6.8, 4% SDS, 0.2% Bromophenol blue, 20% glycerol, 200 mM β mercaptoethanol) and boiled at 100°C for 10 minutes. The samples were cooled to room temperature and 50 μL of distilled water and 10 μL of Proteinase K solution were added and incubated at 60°C for 2 hours. After the incubation, 10 µL of LPS preparation was separated using 14% SDS-PAGE gel. After electrophoresis, the polyacrylamide gel was kept in fixative Solution - A (containing 30% Methanol and 10% Acetic acid in water) overnight. On the next day, the gel was washed with distilled water once by shaking on a rocker for 5 minutes and followed by incubation in Solution - B solution (0.7% periodic acid in fixative A) for 10 minutes. After washing four times with water for (5 minutes each), the gel was incubated in silver stain solution (2.5 mL of 20% silver nitrate + 14 mL of sodium hydroxide solution (1M) + 1 mL of ammonium hydroxide solution + 57 mL of distilled water) for 10 minutes followed by washing (four times) with distilled water. The developing solution (0.05% formaldehyde in 0.28 M sodium carbonate) was added and the gels were developed till the bands turned deep brown. The reaction was stopped by the addition of distilled water.

### 2.8 Bacterial outer membrane (OMP) isolation

*Escherichia coli* cells (10^8^ CFU/ mL) were treated with 1000 µM CuNPs and CuSO_4_ for 4 hours. After the treatment, the cell pellets were re-suspended in 500 µl of 10 mM Tris - pH 7.2 and 5µl of 1 mM PMSF. The mixture was sonicated for 15 seconds 6 times followed by centrifugation at 6000 rpm for 5 minutes at 4°C. The supernatant was centrifuged at 10,500 g for 1 hour at 4°C. The resultant pellet was re-suspended in 50 µl of 10 mM Tris pH 8.0. To this mixture 500 µl of HEPES buffer-pH 7.4 and 500 µl of 1%N-Lauryll sarcosine sodium solution was added and incubated at room temperature for 15 minutes. It was followed by centrifugation at 10,500 g for 1 hour at 4°C. The pellet (OMP) was washed with HEPES buffer and re-suspended in 50 µl of 10% SDS (Sengupta et al. 1992). The concentration of OMP was estimated using the Bamburg method (Minamide and Bamburg 1990). To study the OMP profile, 8 µg of these OMP preparations were separated using 10% SDS-PAGE and stained by using Coomassie Brilliant Blue to visualize the protein bands.

### 2.9 MALDI-TOF-TOF analysis of outer membrane proteins

The gel pieces containing desired proteins were cut carefully, digested with trypsin, and analyzed for the peptide fragments using Bruker Ultraflex MALDI TOF/TOF in the proteomics facility, Indian Institute of Science, Bangalore. The MALDI spectra obtained were loaded into Bruker Daltonics Biotools 3.2 and compared with the *Escherichia coli* database in Swissprot using ‘MS/MS ion search’ in Mascot Software (Matrix Sciences, UK).

### 2.10 Inductively coupled plasma mass spectroscopy (ICPMS) analysis of accumulated copper inside bacterial cells

*Escherichia coli* cells (10^8^ CFU/mL) in nutrient broth were treated with 250, 500 µM of CuNPs and CuSO_4_ for 4 hours. A bacterial culture without any treatment was considered as a control sample. All the samples were done in triplicates. After the treatment, cells were washed twice with phosphate buffered saline with 50 mM glycine (PBSG)) buffer containing 20 µM EDTA to remove the copper present on the surface of the cells. The wet weight of the cell pellet was measured and they were transferred into clean glass tubes by resuspending them in 30 µL of Milli-Q water. To this suspension, 500 µL of 65% HNO_3_ was added and incubated at 70^0^C for 3 hours. After the incubation, the sample was diluted to a 2% final concentration of HNO_3_. The diluted sample was filtered by using 0.45-micron filter to remove any suspended particles and stored at 4^0^C till analysis. On the day of analysis, a standard curve was first made by analyzing various concentrations of copper standard (10, 50, 100, 250, 500 ppb). All the experimental samples were analyzed with XSERIES ICP-MS, Thermo Scientific system in triplicates (Santo et al. 2011).

### 2.11 Statistical analysis

In the case of the antimicrobial study and ICP-MS analysis, statistical significance was calculated using an unpaired t-test. In the case of the measurement of the surface area of the bacteria, the Mann-Whitney U-Test was employed to understand the significance.

## 3. Results

### 3.1 Nanoparticle characterization

TEM analysis confirmed the presence of nano sized particles in the suspension of CuNPs prepared in the laboratory with an average particle size ∼ 4 nm (**Figure 1**).

**Fig. 1.**
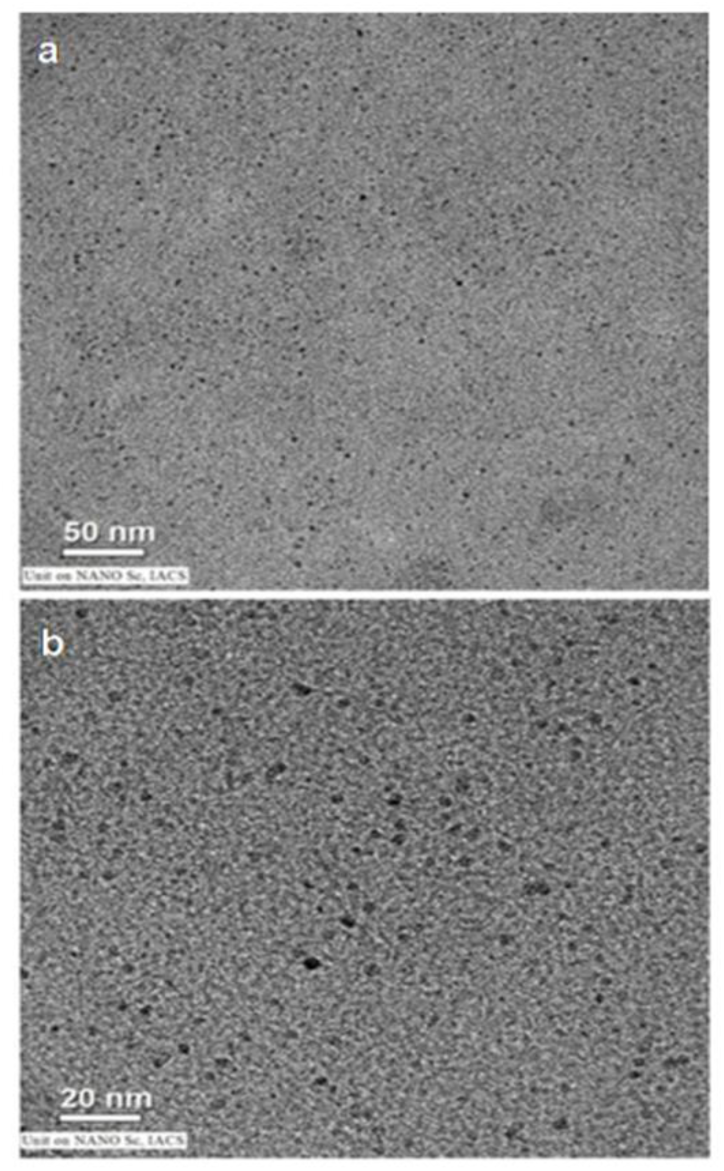
Characterization of copper nanoparticles. Transmission Electron Microscope images of copper nanoparticles at different magnifications.

### 3.2 Antibacterial action of copper nanoparticle vs ionic copper

An antimicrobial study to compare the effect of nano and ionic copper showed that nano copper induced a better bactericidal potential than CuSO_4_ in all concentrations tested and a dose dependency was observed (**Figure 2**) In lower dose (250 µM) copper nanoparticle induced a 54% reduction in CFU compared to 37% caused by ionic copper. With the increase in the dose (500 µM), increased cell death was observed in the case of copper nanoparticles accounting for 89% cell death while ionic copper could only induce 52% cell death. When we increased the dose to 1000 µM, hardly any cells could be seen in the presence of copper nanoparticles however nearly 35% of cells were still viable against ionic copper.

**Fig. 2.**
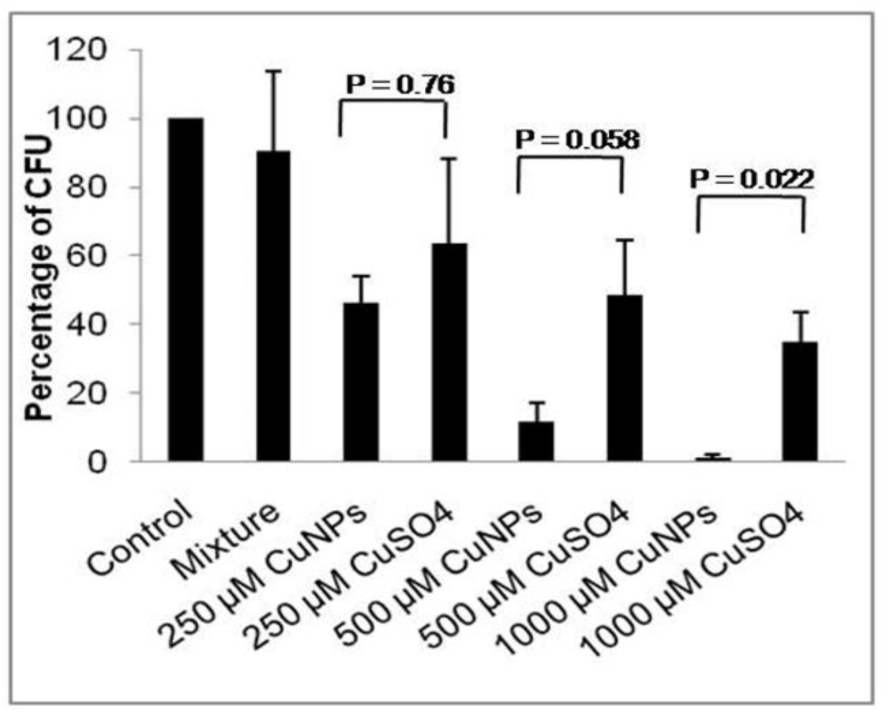
Bactericidal potential of CuNPs/CuSO_4_ on *Escherichia coli*. CuNPs/ CuSO_4_ induces dose dependent bactericidal effect on *Escherichia coli*. Data points are represented as mean ± standard error from three independent experiments.

### 3.3 *In vivo* nucleic acid degradation

To understand the *in vivo* effect of ionic and nano copper in bacteria, the nucleic acid pool from the bacterial cells was isolated after the CuNPs / CuSO_4_ treatment. A dose dependent increase in the nucleic acid (both DNA and ribosomal RNA) degradation by CuNPs was observed (**Figure 3**). Interestingly, total degradation of ribosomal RNA pool was observed for 750 µM and higher concentrations of CuNPs clearly indicating that higher doses of CuNPs are more effective. In the case of CuSO_4_ treatment, there was no nucleic acid degradation was observed till the 1000 µM dose. Only 1500 µM of CuSO_4_ showed both genomic DNA and ribosomal RNA degradation.

**Fig. 3.**
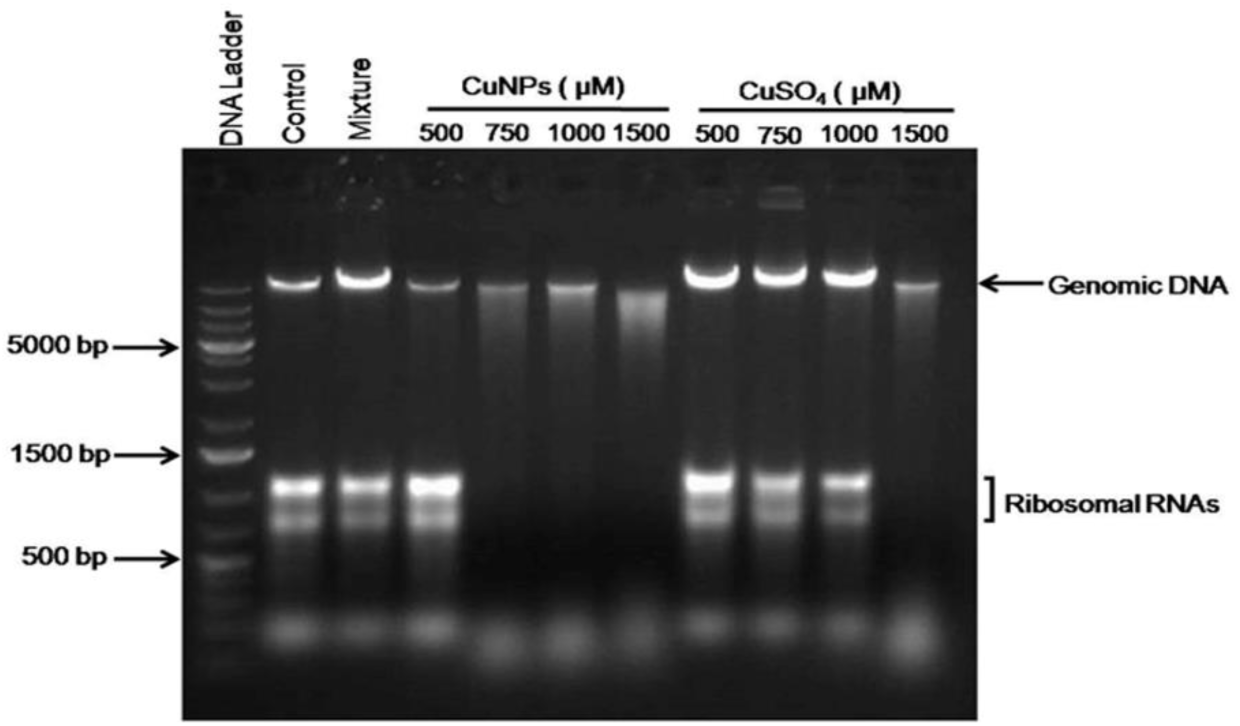
Comparison of nucleic acid degradation potential of CuNP/CuSO_4_. Representative image of nucleic acid degradation profile of *Escherichia coli* cells treated with various concentrations of CuNPs/CuSO_4_

### 3.4 Reactive Oxygen Species generation

Fluorescence microscopic studies of E. coli cells stained with DCFDA were employed to understand the involvement of ROS in the antibacterial activity induced by CuNPs. Results (**Figure 4**) showed that the CuNPs can induce the generation of ROS in a greater number of bacterial cells compared to CuSO4 in the presence of 1000 µM CuNP. Control cells (without any treatment) showed a deficient number of fluorescent cells indicating the absence of induction of ROS generation. Hydrogen peroxide treated cells (as positive control) showed intense fluorescence as expected.

**Fig. 4.**
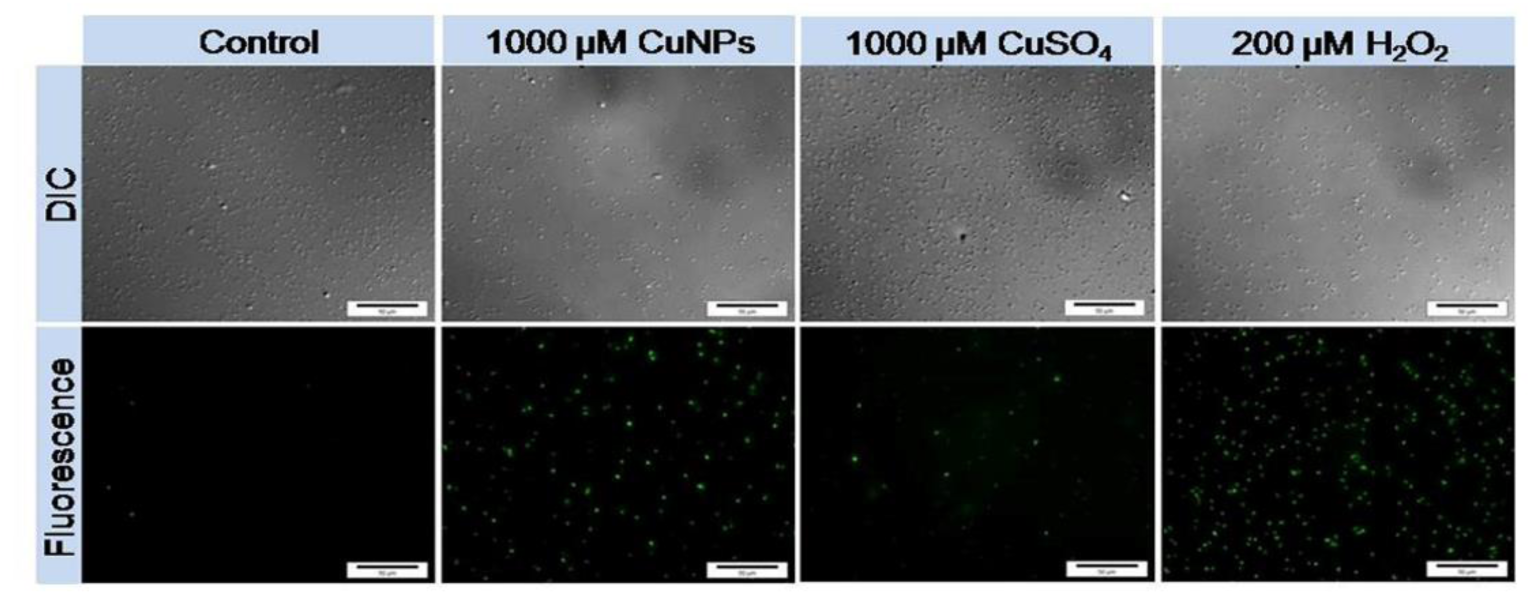
ROS generation in the presence of CuNPs/CuSO_4_. Representative images of ROS detection in *Escherichia coli* cells after treatment with CuNPs/CuSO_4_. (Magnification 60 X)

### 3.5 Bacterial membrane damaging action

The scanning electron microscopic (SEM) analysis showed a striking difference in surface morphology between the nano and ionic copper treated Escherichia coli cells. When treated with 1000 µM of CuNPs, *Escherichia coli* cells appeared to have the presence of membrane pits which were completely absent in 1000 µM CuSO_4_ treated cells (**Figure 5a**). Also, a significant increase in the surface area was observed in the case of CuNPs treated cells compared to the control or ionic copper treated *Escherichia coli* cells (**Figure 5b**).

**Fig. 5.**
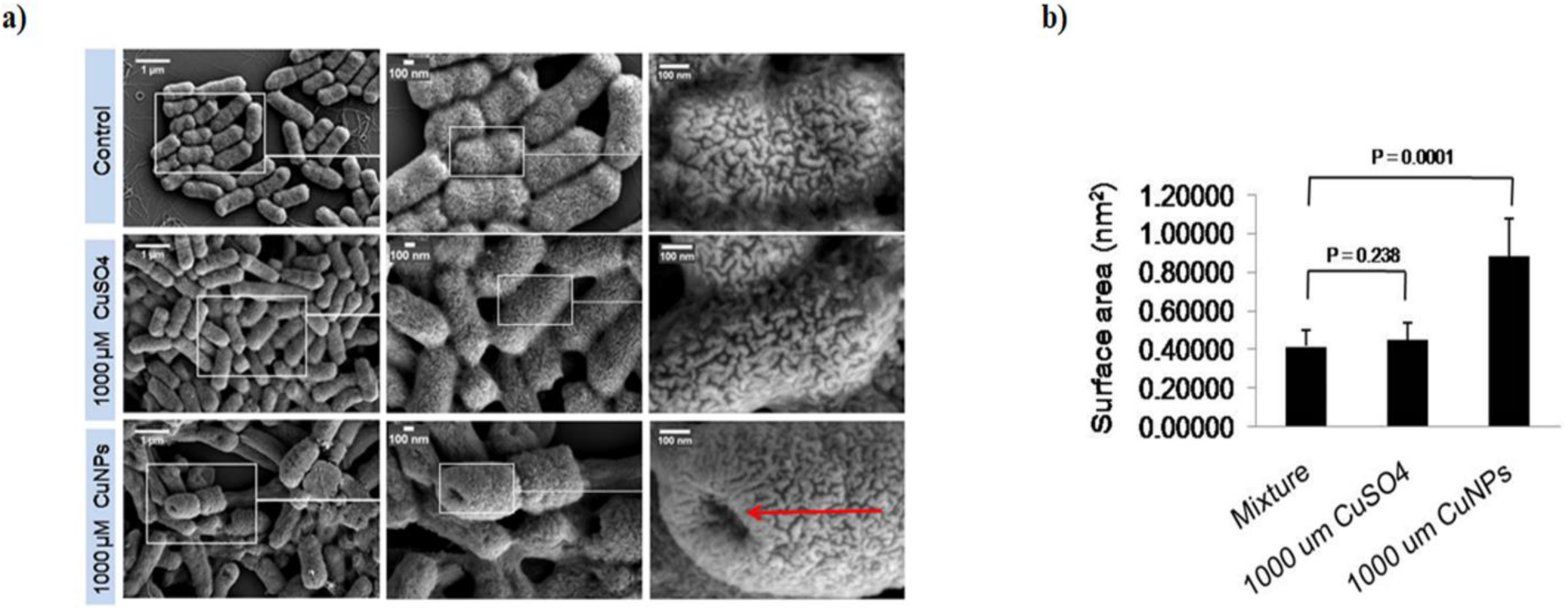
Scanning Electron Microscopic Analysis. **(a)** Representative images of SEM analysis of *Escherichia coli* cells after treating with 1000 µM CuNPs and 1000 µM CuSO_4_ **(b)**, Comparison of surface area of *Escherichia coli* cells treated with 1000 µM CuSO_4_ and 1000 µM CuNPs. Data points for area measurement correspond to mean ± standard error from 10 independent measurements. Red arrow indicates the presence of membrane pit.

### 3.6 Lipopolysaccharide profile

Cell surface lipopolysaccharide (LPS) in Gram-negative bacteria acts as a protective layer against various antibacterial agents like long-chain fatty acids, antimicrobial peptides, etc (Papo and Shai 2005; Sheu and Freese 1973). However, LPS profile (O-antigens) analysis of the *Escherichia coli* cells treated with CuNPs or CuSO_4_ showed no significant difference (**Figure 6**).

**Fig. 6.**
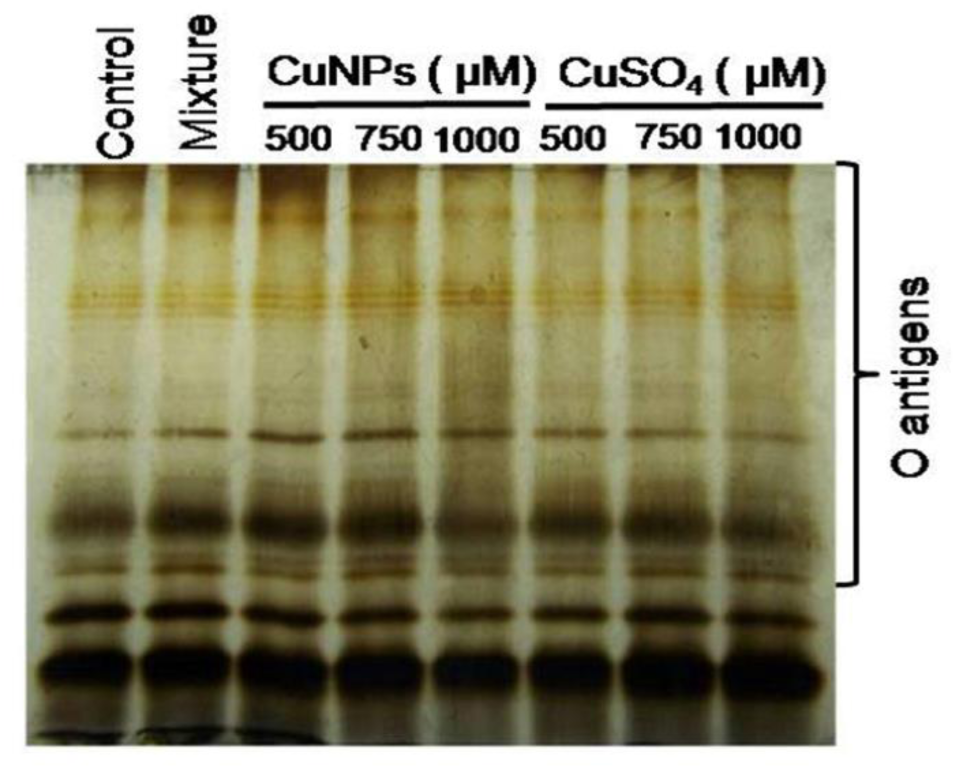
SDS-PAGE analysis of lipopolysaccharide. Representative image of lipopolysaccharide profile of *Escherichia coli* cells treated with various concentrations of CuNPs/ CuSO_4_

### 3.7 Outer membrane protein profile

Analysis of the outer membrane protein (OMP) profiles of *Escherichia coli* cells revealed striking differences in its composition in response to interaction with CuNPs compared to untreated cells. Interestingly, it showed a dose dependent down-regulation of maltoporin (LamB) and outer membrane protein A (OmpA) along with the up-regulation/ translocation of elongation factor Tu (EF-Tu) in CuNP treated cells (**Figure 7**). CuSO4 could also exert a similar effect but at higher concentrations compared to CuNPs. The up- and down-regulated proteins were identified by mass-spectrometry and the **Table 1** represents the Mascot server predicted proteins with MS-MS score.

**Table 1.** Mascot server predicted proteins with MS MS score.

| **Band** | **Molecular weight (kDa)** | **Peptide mass fingerprinting prediction** | **MS MS score** |
| --- | --- | --- | --- |
| **P1** | 47 | Maltoporin (LamB) | 505 |
| **P2** | 44 | Elongation factor Tu | 204 |
| **P3** | 34 | Outer membrane protein A | 46 |
| **P4** | 21 | Alkyl hydro peroxide reductase subunit C | 34 (MS score) |

**Fig. 7.**
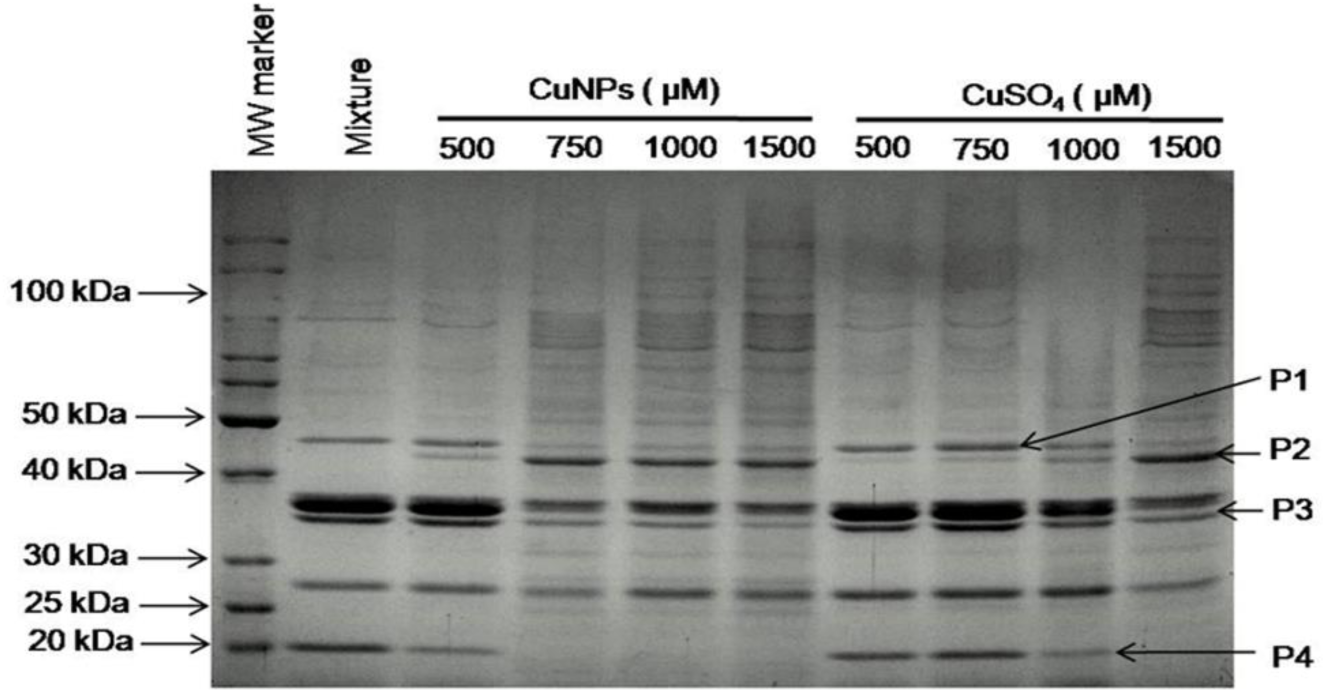
Outer membrane protein analysis. Representative image of SDS-PAGE analysis of outer membrane proteins isolated from *Escherichia coli* cells after treatment with CuNPs/ CuSO_4_

### 3.8 Entry of copper in to bacterial cells

ICP-MS analysis showed a significantly increased (2.42 folds) presence of copper in CuNPs treated E. coli cells than the CuSO_4_ treated cells at 250µM concentration. At 1000µM concentration, CuSO_4_ treated cells had higher intracellular copper levels in comparison to nano copper treated cells (**Figure 8**). This unusual observation may have been due to the fact that at 1000µM concentration of CuNPs, the bacteria undergo membrane damage by pit formation as seen in our scanning electron microscopy results **(Figure 5)**, thereby rendering the bacteria unable to retain CuNPS inside their cells.

**Fig. 8.**
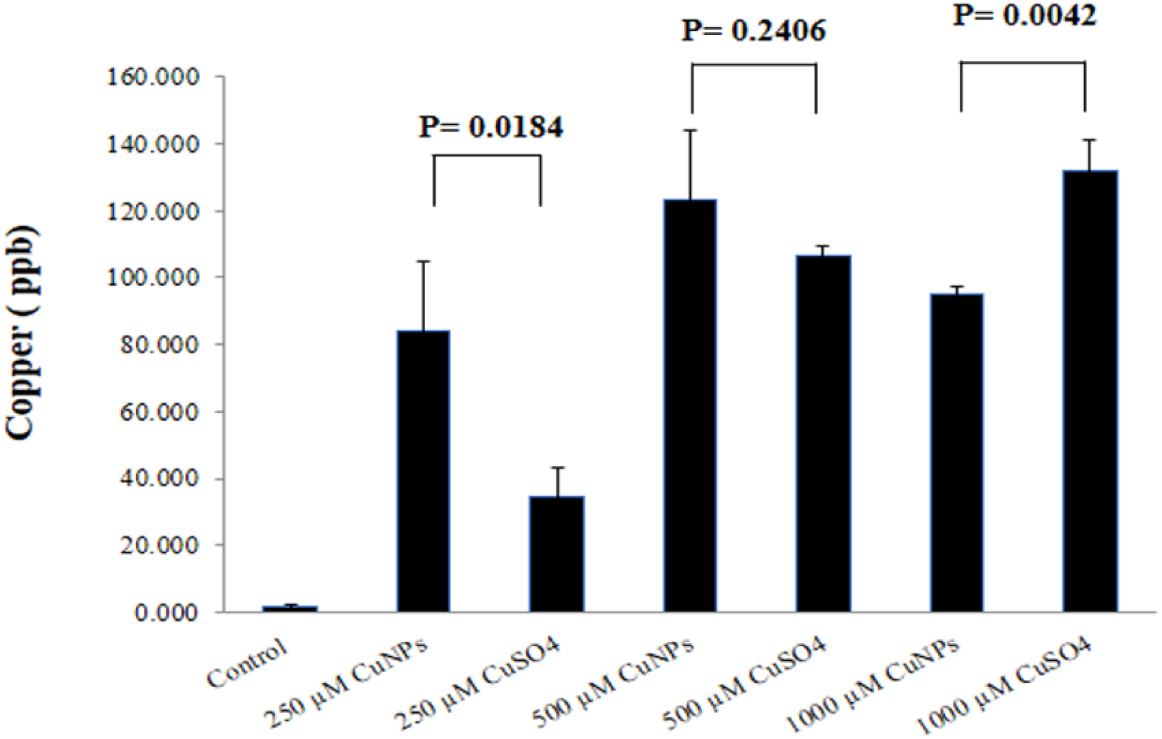
ICPMS analysis. Amount of copper present in *Escherichia coli* cells, after the treatment with various concentrations of CuNPs/ CuSO_4_. (Each data point represents the mean ± standard error from three independent samples)

## 4. Discussion

Understanding the difference in the mechanism of antibacterial action of ionic and nano copper is important to ascertain the advantage of using nano copper over ionic copper. This study comprehensively addresses the difference in the nano and ionic copper mediated cell death in bacteria using *Escherichia coli* as a model organism. This study shows the *in vivo* DNA degradation potential of CuNPs in comparison to ionic copper in *Escherichia coli*. It was reported that copper surfaces can cause a rapid bactericidal action associated with the degradation of both genomic and plasmid DNA in pathogenic *Enterococci* (Warnes et al. 2010). It was also observed that in the case of Gram-negative bacteria like *Escherichia coli* and *Salmonella*, the DNA destruction rate is slow (Warnes et al. 2012,). Previous reports showed that CuNPs alone can cause DNA degradation *in vitro*, mediated by the generation of singlet oxygen (Jose et al. 2011). But in the case of bacteria, there are number of enzymes specialized in the defense against the detrimental effects of reactive oxygen species (ROS) (Storz et al. 1990). Therefore, to understand the *in vivo* effect of ionic and nano copper in bacteria, the nucleic acid pool from the bacterial cells was isolated after the CuNPs / CuSO_4_ treatment. A dose dependent increase in the nucleic acid (both DNA and ribosomal RNA) degradation by CuNPs was observed (**Figure 3**). The *in vivo* nucleic acid degradation activity shown by CuNPs at lower concentrations even in the presence of efficient free radical scavenging systems in bacteria is note-worthy as extensive nucleic acid degradation activity will prevent the bacteria from acquiring any resistance towards CuNPs toxicity.

To understand the involvement of ROS in the antibacterial activity induced by CuNPs, fluorescence microscopic studies of *E. coli* cells stained with DCFDA were employed. Results (**Figure 4**) showed that the CuNPs can induce generation ROS in a greater number of bacterial cells compared to CuSO4 when the same concentrations were used. Control cells (without any treatment) showed very low number of fluorescent cells indicating the absence of induction of ROS generation. The ability of copper to generate ROS, mediated by the Fenton reaction in biological systems well known (Cabiscol et al. 2000). Copper can cause the generation of superoxide and hydroxyl radicals by accepting or donating electrons and cycle between cuprous and cupric forms (O’Gorman and Humphreys 2012). The present study showed that CuNPs could induce a greater extent of oxidative stress in bacteria than CuSO_4_.

The scanning electron microscopic (SEM) analysis showed a striking difference between the nano and ionic copper treated *Escherichia coli* cells in their surface morphology. When treated with 1000 µM of CuNPs, *Escherichia coli* cells appeared to have the presence of membrane pits which were completely absent in 1000 µM CuSO_4_ treated cells (**Figure 5a**). Also, a significant increase in the surface area was observed in the case of CuNPs treated cells compared to the control or ionic copper treated *Escherichia coli* cells (**Figure 5b**). Membrane damage caused by CuNPs is not a novel phenomenon as there are reports that showed copper nanoparticle mediated membrane pit formation in bacteria (Li et al. 2013; Raffi et al. 2010). However there are no clear explanations, based on experiments, available for such membrane pit formation by CuNPs. It was reported before that silver nanoparticles can cause such membrane pit formation in *Escherichia coli* cells (Sondi and Salopek-Sondi, 2004). The most plausible explanation for such membrane damage caused by metal nanoparticles could be based on the interaction of such nanoparticles with membrane proteins, mainly those proteins which contain sulfhydryl (–S– H) groups to form R–S–S–R bonds, thus inactivating their function (Cho et al. 2005).

Cell surface lipopolysaccharide (LPS) in Gram negative bacteria acts as a protective layer against various antibacterial agents like long-chain fatty acids, antimicrobial peptides, etc (Papo and Shai 2005; Sheu and Freese 1973). LPS profile (O-antigens) analysis of the *Escherichia coli* cells treated with CuNPs or CuSO_4_ showed no significant difference. This result showed that the antibacterial activity of either CuNPs or CuSO_4_ was not mediated through the alteration of the LPS layer (**Figure 6**).

Analysis of outer membrane protein (OMP) profiles of *Escherichia coli* cells revealed striking differences in its composition in response to interaction with CuNPs compared to untreated cells. Interestingly, it showed a dose dependent down-regulation of maltoporin (LamB) and outer membrane protein A (OmpA) along with the up-regulation/ translocation of elongation factor Tu (EF-Tu) in CuNP treated cells (**Figure 7**). CuSO4 could also exert a similar effect but at higher concentrations compared to CuNPs. The up- and down-regulated proteins were identified by mass-spectrometry and the **Table 1** represents the Mascot server predicted proteins with MS-MS score. EF-Tu is one of the prokaryotic elongation factors that help in the translation of proteins by recruiting aminoacyl-tRNAs to fit into ribosomes during the process of translation. The first report about the membrane association of EF-Tu was described by Jacobson and Rosenbusch (Jacobson and Rosenbusch 1976). Later, many studies confirmed that this translocation is associated with osmotic shock (Berrier et al. 2000; Lunn and Pigiet 1982; Vazquez-Laslop et al. 2001). The present study also shows that CuNPs induced an osmotic shock by creating membrane pits leading to the loss of membrane integrity of the bacterial cells and that may be the probable reason for the translocation of EF-Tu on to the membrane fraction. OmpA is primarily involved in the stability of bacterial membrane (Wang 2002). It connects the outer membrane with the peptidoglycan layer and helps the cell to maintain its shape in harsh conditions. The downregulation of OmpA protein by CuNPs supports the role of CuNPs in destabilizing the bacterial membrane. LamB porins are required by *Escherichia coli* for the diffusion of maltodextrins into the cells (Boos and Shuman 1998). Downregulation of LamB is mainly seen in different antibiotic-resistant strains of bacteria (Lin et al. 2010). However, the role or involvement of LamB in CuNPs treated bacterial cells was not explored further in the present study. Interestingly, alkyl hydroperoxide reductase subunit C (AhpC) also got downregulated due to CuNP treatment (but the MS score was not significant). A similar downregulation of AhpC was seen in *Escherichia coli* cells treated with gold nanoparticles (Cui et al. 2012).

ICP-MS analysis showed a significantly increased (2.42 folds) presence of copper in CuNPs treated E. coli cells as compared to the CuSO_4_ treated cells at 250µM concentration and 500 µM concentration showed 1.15 fold increase in copper concentration in CuNPs compared to CuSO_4_ whereas at 1000 µM concentration, there was no significant change observed in the copper level between the nano and ionic copper treated cells (**Figure 8**) presumably due to the inability to retain the nano coppers as a result of membrane pit formation (**Figure 5**). These results indicate better internalization of CuNPs in *E. coli* cells compared to CuSO4 and pointing towards the ‘Trojan horse’ mechanism for antibacterial activity of CuNPs at low concentrations.

Based on the results, it can be concluded that the nano nature of copper has a major advantage over ionic copper in inducing bactericidal activity. The increased sensitivity of *Escherichia coli* cells towards CuNPs can be ascribed to the relatively higher ability of CuNPs to trigger *in vivo* nucleic acid degradation, ROS generation, and membrane damaging action. The changes in the outer membrane protein profile, most likely mediated through osmotic shock, were more evident in CuNPs treated cells. These observations also point towards a potent interaction of CuNPs with bacterial membranes. ICP-MS studies revealed better entry of CuNPs into the bacterial cells, thus indicating the ‘Trojan Horse’ mechanism for CuNPs mediated antibacterial activity. In summary, CuNPs can be considered as a better candidate for fighting against nosocomial infections. But, the issues like oxidation of CuNPs and nanoparticles leaching into the environment should be addressed properly before large-scale applications.

## Statement and Declaration

### Author contribution

SS prepared the nanoparticle. GPJ performed all the experiments. GPJ, SKS, TKS analyzed the data and wrote the manuscript. TKS and SM conceived the study and reviewed the manuscript.

### Availability of Data and Materials

In this study, all data generated or analyzed are included in the article.

### Ethical Approval

Not applicable.

### Consent to Participate

All authors had consented to participate in the study.

### Consent for Publication

All authors have given consent for publication.

## Acknowledgments

Gregor P. Jose and Subhankar Santra thank the Council of Scientific and Industrial Research, Government of India for research fellowships. Saurav Kumar Saha would like to acknowledge Indian Institute of Science Education and Research, Kolkata for providing fellowship. The authors want to acknowledge the TEM facility of the Indian Association for the Cultivation of Science, India. Swadhin K Mandal thanks the Department of Science and Technology, Government of India for financial support. The authors wish to thank the Indian Institute of Science Education and Research (IISER) Kolkata for providing infrastructure, SEM facility and ICP-MS facilities of IISER Kolkata. The authors acknowledge the financial support provided by IISER Kolkata.

